# Investigating nanostructure and -mechanics of contracting actin stress fibers by scanning ion conductance microscopy

**DOI:** 10.64898/2026.07.01.735957

**Authors:** Yanjun Zhang, Yasufumi Takahashi, You-Rong Lin, Andrew Shevchuk, Yuri Korchev, Clemens M. Franz

**Affiliations:** WPI Nano Life Science Institute, Kanazawa University, Ishikawa 920-1192, Japan; Department of Electrical Engineering, Graduate School of Engineering, Nagoya University, Aichi 464-8601, Japan; Research Institute for Quantum and Chemical Innovation, Institutes of Innovation for Future Society, Nagoya University, Furo-cho, Chikusa-ku, Nagoya 464-8601, Japan; Exploratory Research Center on Life and Living Systems (ExCELLS), National Institutes of Natural Sciences, 5-1 Higashiyama, Myodaiji, Okazaki, Aichi 444-8787, Japan; Institute for Molecular Science (IMS), National Institutes of Natural Sciences, 5-1 Higashiyama, Myodaiji, Okazaki 444-8787, Japan; Department of Metabolism, Digestion and Reproduction, Imperial College London, London W12 0NN, United Kingdom

**Keywords:** SICM, SPM, actin stress fibers, cell mechanics, actomyosin contractility

## Abstract

Scanning ion conductance microscopy (SICM) provides gentle, non-contact cell surface imaging, but it has not been used to investigate intracellular structures because the plasma membrane restricts nanopipette access. Here, we combined SICM with microsonication-based cell de-roofing to expose intracellular actin stress fibers (SFs) in U2OS cells for nanotopographical and -mechanical characterization. Importantly, the de-roofing conditions preserved actomyosin contractility, allowing analysis of SF structural and biomechanical changes during ATP-induced contraction. Resting SFs displayed an average height of 203±38 nm and width of 357±73 nm, and a complex surface architecture characterized by regularly spaced long-range height modulations (∼500 nm periodicity; Wq ∼25 nm) and smaller irregular corrugations (Ra ∼19.2 nm). ATP stimulation reduced SF height and width by ∼39% and ∼15%, respectively, while largely preserving surface corrugation patterns. During contraction, some SFs separated into two longitudinal strands. High-resolution SICM imaging also revealed filamentous crosslinks mechanically coupling neighboring SFs, and nanomechanical measurements demonstrated local stiffening during contraction. These findings provide new insight into the structural and mechanical regulation of SF contraction and highlight the potential of SICM combined with cell de-roofing as a powerful platform for studying dynamic intracellular processes at nanometer resolution.

## Introduction

Stress fibers (SFs) are prominent intracellular bundles of actin filaments found in a variety of non-muscle cells. Within SFs individual actin filaments are arrayed in parallel and crosslinked by several actin-binding proteins, including α-actinin, zyxin, and filamin^1^. Importantly, SFs contain non-muscle myosin II motors, and myosin-driven antiparallel sliding of actin filaments leads to strong SF contraction, and consequently, increased intracellular contractility^2^. Because the ends of SFs are linked via integrin-containing focal adhesion (FA) complexes to the extracellular environment, SF contraction enables cells to exert forces against the extracellular surrounding, which is crucial for maintaining intracellular tension and for resisting external deformation. SF-based contractility underlies numerous cellular processes, such as cell-matrix adhesion, migration, wound healing, and mechanosensing, and changes in the direction or magnitude of SF-dependent forces can contribute to diseases such as atherosclerosis, osteoporosis, and cancer^3,4^.

Intracellular signaling pathways regulating myosin II-dependent SF contraction have been elucidated in great detail^5^. Likewise, SF and FA ultrastructure has been investigated extensively by scanning- and transmission-electron microscopy (EM)^6,7^, cryo-EM^8^, or helium ion microscopy^9^. However, these microscopy techniques only capture static snapshots of different structural states of SFs and thus cannot readily reveal the dynamic structural and mechanical changes that drive SF contraction and force generation. Optical superresolution microscopy has provided valuable insight into the dynamic organization of myosin II motor assemblies within SFs^10^. However, it remains challenging to resolve the densely packed individual actin filaments and to obtain direct mechanical information. Therefore, microscopy techniques capable of operating under physiological conditions and that simultaneously provide nanoscale structural and mechanical information are required. Scanning ion conductance microscopy (SICM) is a non-contact scanning probe microscopy (SPM) variant that enables real-time acquisition of nanometer resolution topographical images of biological samples under physiological conditions^11–13^. SICM reconstructs sample topography by monitoring the decrease in ionic current flowing through a glass nanopipette as it approaches the sample surface. As a non-contact imaging technique, SICM applies minimal mechanical perturbation during scanning, making it particularly suitable for visualizing fragile biological structures, such as microvilli, microridges, and other highly dynamic membrane protrusions and corrugations on living cells^14,15^.

The development of hopping probe mode SICM (HPICM) further reduces, or effectively eliminates, residual lateral forces during scanning^16^. In HPICM, the nanopipette is first withdrawn to a position well above the sample surface and then approaches the sample vertically from above, thereby avoiding lateral collisions between the probe and the sample. In this manner, even extremely fragile cellular structures, such as intricate dendritic networks, can be imaged at high-resolution over extended periods without incurring detectable damage^16^. Importantly, SICM also provides the ability to quantify mechanical properties of living cells^17^, for instance related to fragile and highly dynamic membrane protrusion on the surface of living cancer cells^18^. However, as a surface scanning technique, SICM investigations are generally limited to processes associated with the plasma membrane, such as membrane ruffles and microvilli^15^, and endocytosis events^19^, while intracellular structures remain inaccessible. Methods that enable direct access of the SICM nanoprobe to specific intracellular structures are therefore highly desirable. One promising approach is the gentle removal of the plasma membrane while preserving the structural and functional integrity of the underlying intracellular components.

Cell de-roofing using detergent extraction or a series of controlled ultrasonic bursts in buffers emulating intracellular conditions^20–24^ can expose intracellular structures in their native state for subsequent nanoprobe exploration^25^. Although cell de-roofing provides the opportunity to investigate intracellular structures by different SPM techniques, actin structures in de-roofed cells have thus been examined exclusively by atomic force microscopy (AFM), either in their native state^26^ or following chemical fixation^22,23^ to minimize sample deformation during scanning. With careful adjustment of lateral and vertical scan forces, AFM can resolve intricate molecular assemblies in de-roofed cells with nanometer resolution^25^. However, imaging large, highly corrugated intracellular landscapes and fragile organelles remains challenging and often requires chemical fixation to stabilize the sample during scanning^27^. Given its non-contact and minimally invasive imaging characteristics, SICM may therefore be more suitable than AFM for investigating large intracellular protein complexes and organelles, but so far it has not yet been applied to the direct imaging of intracellular structures. Here we combined cell de-roofing, SICM, and fluorescence microscopy to investigate the ultrastructure and mechanical properties of actin SFs in de-roofed U2OS osteosarcoma cells. De-roofing was performed under conditions that preserved SF contractility in response to external ATP stimulation^28,29^, enabling direct visualization of structural and mechanical changes associated with SF contraction. Besides providing new insight into SF contraction that cannot be obtained with other techniques, the combination of SICM and cell de-roofing may serve as a powerful platform for investigating other dynamic intracellular processes in the future.

## Results

### Correlative SICM and fluorescence imaging of the exposed actin SFs

Among SPM techniques, the non-contact nature of SICM makes it uniquely suited for visualizing fragile cellular structures, including dynamic microvilli or membrane ruffles on the surface of living cells^14,18,19^. In principle, SICM should also be well-suited for imaging delicate intracellular structures and organelles in de-roofed cells, which often require chemical fixation to achieve high-resolution imaging by AFM^22,23^. Here, we explore the application of SICM for high-resolution imaging of intracellular actin cytoskeleton structures. To gain nanopipette access to the cell interior, we first removed the apical plasma membrane from U2OS cells using a modified microsonication-based cell de-roofing method^22^. This procedure removes most intracellular organelles and the nucleus while preserving the basal plasma membrane together with the associated cortical actin cytoskeleton and SFs, including their focal adhesion (FA)-associated termini. By adjusting the microsonication strength, cells could be partially (Suppl. Fig. S1B) or fully deroofed (Suppl. Fig. S2), progressively revealing intracellular structures and organelles. Using a U2OS cell line stably expressing actin-GFP allowed us to identify SFs and to compare their arrangement before (Fig. 1A) and after de-roofing (Fig. 1B) by fluorescence microscopy. Overall, SFs were well-preserved after de-roofing, although some fibers were disrupted during the procedure, typically in central regions where they had previously been connected to the perinuclear matrix^30^ (Fig. 1A). Fluorescence image contrast was enhanced in de-roofed samples due to the removal of cortical actin structures associated with the apical plasma membrane and loss of diffuse cytosolic G-actin fluorescence signals. Next we imaged de-roofed cells by SICM using 20 x 20 µm^2^ overview scans with 256 x 256 pixel resolution, yielding a pixel size <80 nm. SICM topographs showed a similar SF arrangement as the fluorescence images but provided substantially greater structural detail, including the apparent fanning out of individual actin filaments at the distal FA end, occasional crossovers of SFs, and abundant intricate interfibrillar connections (Fig. 1C). Furthermore, the basal plasma membrane was detected ∼10 nm above the glass substrate, confirming that the de-roofing procedure preserves the intact basal plasma membrane together with its associated cytoskeletal structures. We subsequently re-scanned 10 × 10 µm^2^ subregions of the de-roofed cell sample at higher spatial resolution (256 x 256 pixel grids, pixel size <40 nm, Fig. 1D and 1E). Again, SICM and fluorescence images showed good overall agreement, but the SICM images contained greater ultrastructural detail than the optical images, revealing a series of globular structures distributed along the SFs as well as extensive interfibrillar bridges. We estimated the lateral (*X*/*Y)* resolution of the SICM images to be ∼50 nm, consistent with the nanopipette tip diameter used in the present study and our previous high-resolution SICM investigations^31^. Although the SICM Z-positioning system can achieve high precision due to sensitive ion current feedback and piezoelectric control, the experimentally demonstrated vertical resolutions in biological imaging are typically on the order of 10 nm^16,32^. These results demonstrate that SICM can visualize intracellular structures in de-roofed cells with near-molecular resolution and superior three-dimensional (*X*/*Y*/Z) resolution compared to conventional fluorescence microscopy, as well as higher Z resolution than optical superresolution microscopy^33^.

**Figure 1.**
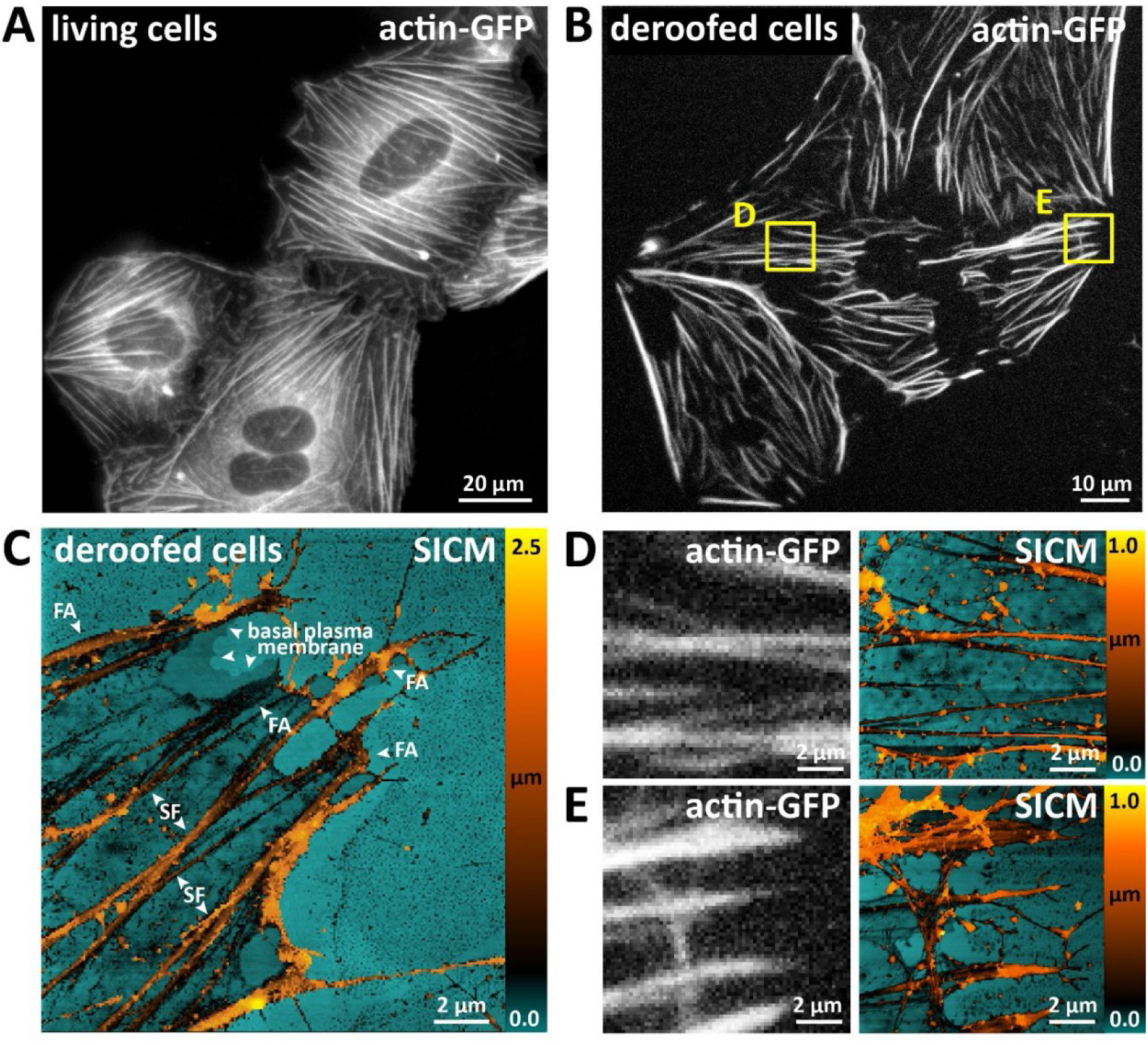
Cell de-roofing for intracellular exploration by SICM. **(A)** Living U2OS cells stably expressing tagGFP-actin at 60x magnification. **(B)** Exposed SFs in deroofed U2OS**-**tagGFP cells. Yellow boxes indicate regions shown at higher magnification in **(D**) and **(E**). **(C)** SICM topographic overview image (20 x 20 µm^2^, 256 x 256 pixel, pixel size ∼78 nm) of a deroofed cell revealing the arrangement of stress fibers (SF), focal adhesions (FA) and the basal plasma membrane. **(D)** and **(E)** Enlarged subregions of the optical image (left) corresponding to the yellow boxes in **(B)** and higher resolution SICM re-scans (10 × 10 µm^2^, 256 x 256 pixel, pixel size ∼40 nm) of the same areas (right).

### Visualizing structural changes in contracting SFs

Several previous AFM studies investigating actin structures in de-roofed cells employed chemical fixation or drying to avoid sample deformation due to lateral forces exerted by the AFM tip during scanning^22,23^. In contrast, the contactless nature of SICM imaging meant that unfixed de-roofed cells samples could be scanned continuously for at least 2 h without noticeable sample deformation or image quality degradation (Suppl. Figure S2). The ability to image unfixed de-roofed cells under physiological conditions provides the opportunity to observe dynamic intracellular molecular processes directly within the native cellular environment. Specifically, the buffer used in de-roofing protocol retains the contractility of the exposed SFs in response to ATP/Mg^2+^ stimulation^28^. We first confirmed SF contractility after de-roofing by simultaneous fluorescence microscopy and optical holotomography imaging. Intracellular ATP concentrations range up to 8 mM^34^. Using repeated doses of a lower ATP/Mg^2+^ concentration (1 mM) applied every 10 min enabled us to induce continuous SF contraction over a time frame of 40 min, after which additional ATP caused no further contraction (Figure 2 and Supplementary Movie 1).

**Figure 2.**
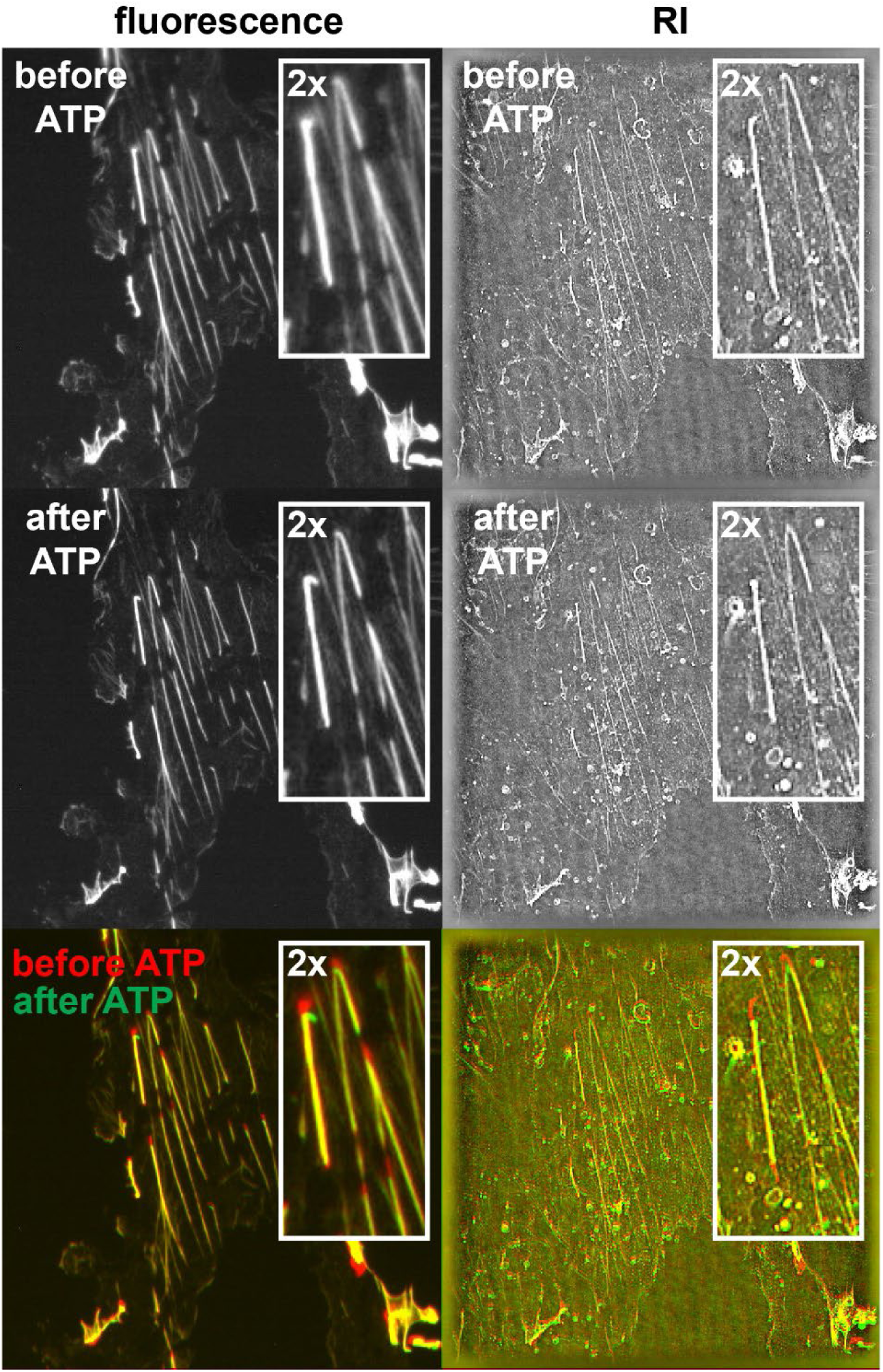
Exposed SFs remain contractile in response to ATP. De-roofed U2OS cells expressing tagGFP-actin were simultaneously imaged by fluorescence (left panels) and 3D holotomography (right panels) before (top) and after stimulation with 1 mM ATP/Mg^2+^. Inserts show twofold magnified views as single channels (top, middle) or a merged image (bottom) to visualize SF contraction. The uncropped image size corresponds to 90 x 90 µm^2^.

Initial SF contraction rates (∼0.2 µm/min) in this *in vitro* assay were comparable to rates (∼0.1 µm/min) observed in living cells^35^. Inducing slow, continuous SF contraction thus enabled us to visualize nanoscale structural changes occurring within contracting SFs by SICM even at limited frame rates. For this we selected areas close to the cell periphery containing several prominent SF/FA termini. To increase image acquisition speed and image resolution, we again reduced the scan region to 10 × 10 µm^2^ while maintaining a 256 x 256 pixel grids, yielding a ∼40 x 40 nm pixel size, close to the expected resolution limit provided by the ∼50 nm nanopipette tip diameter. Operating the nanopipette scanner at the maximum scan speed maintaining reliable sample contouring (∼8.5 µm/sec) yielded an image acquisition time of ∼5 min per frame under these conditions. After 10 min of continuous scanning, we added the first dose of ATP (1 mM). Morphological changes associated with SF retraction became visible in the first image frame after ATP addition (Fig. 3A and 3B, and Supplementary Movie 2). In particular, the FA area at the end of a prominent SF displayed translocation away from the original cell boundary in a sliding movement. The translocation continued over the next 40 min with 1 mM ATP re-stimulation every 10 min, while further ATP stimulation after this time again caused no further FA translocation or SF contraction. While SFs consist of tightly bundled actin filaments^36^, the FA termini are characterized by the gradual fanning out of individual actin filaments into a flattened, fan-shaped arrangement^22^. Interestingly, the SF termini retained this frayed actin filament arrangement during translocation, although with a gradual thinning of actin filament density over time (Fig. 3B), suggesting that basic actin architecture is maintained during FA “sliding”^37^. Tracking local displacements along an individual SF over time revealed that shrinkage occurred predominantly from the SF end (Fig. 2C), while central SF regions showed little or no translocation. This suggested that SFs shrink predominantly by end shortening, rather than by homogenous contraction along their length. Together with SF contraction, ATP stimulation also induced cell membrane retraction (Fig. 3B), consistent with tight mechanically coupling between SFs and the membrane-underlying cortical cytoskeleton. Applying a ridge detection algorithm to the overview scan images of contracting SFs also revealed an intricate, mechanically-coupled network of crosslinks between neighboring fibers (Supplementary Movie 3).

**Figure 3.**
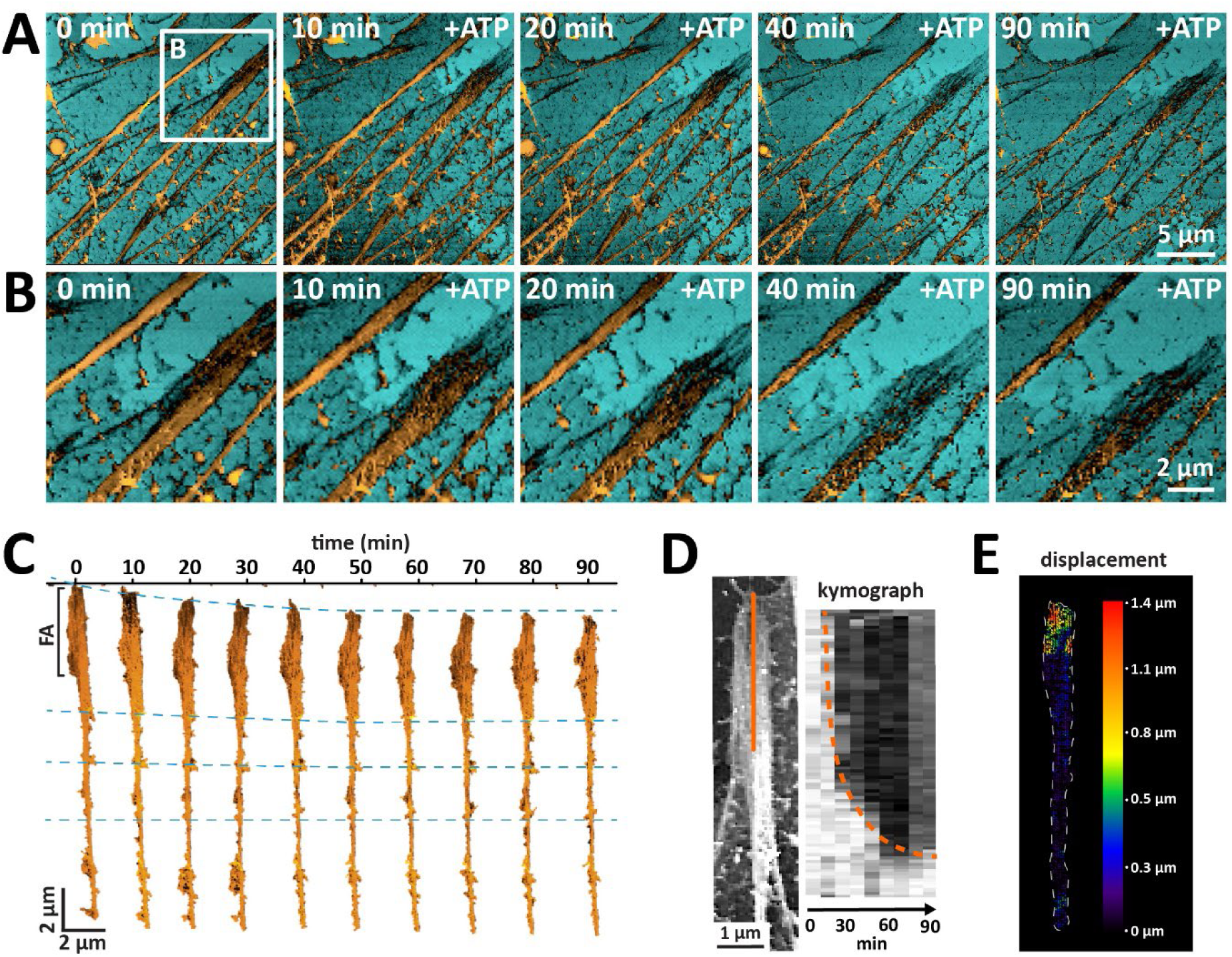
**(A)** Timelapse SICM image series of SFs contracting in response to ATP stimulation. ATP/Mg^2+^ complex was added in individual doses of 1 mM up to a total final concentration of 4 mM. **(B)** Enlarged view from the same timelapse series corresponding to the boxed area in **(A)** depicting the FA region of a contracting SF. The full range of the height scale corresponds to 2 μm. **(C)** An individual SF with associated FA region extracted from the timelapse series. Images were vertically aligned using immobile sample features for reference. Dashed blue lines connect corresponding SF features across timepoints. **(D)** Kymograph (right panel) generated along a cross-section (orange line) at the distal end of a retracting SF. **(E)** Displacement map of a contracting SF, demonstrating that shrinkage occurs predominantly from the SF end.

### Reduction of SF height and width during contraction

The high resolution of the SICM images enabled us to assess additional nanoscale changes to SF structure during ATP-stimulated contraction. 3D reconstructions of SICM topographies demonstrated an overall decrease in SF height after contraction (Fig. 4A and 4B). A representative SICM topographic image from another time-lapse series likewise showed ATP-induced height and width contraction of SFs (Fig. 4C). Cross sections taken along (Fig. 4D) or across (Fig. 4E) contracting SFs demonstrated reductions in SF height (Fig. 4F) from 203 ± 38 to 124 ± 41 nm (mean ± SD, ∼39% reduction) and width from 357 ± 73 to 302 ± 61 nm (∼15% reduction). SICM records sample height with nanometer precision, but lateral sample dimensions are typically inflated due to convolution effect with the nanopipette apex^38^. However, edge effects remain constant, and the net mean lateral fibril shrinkage (357 nm – 302 nm = 55 nm) therefore likely provides an accurate measure of true fiber width reduction. Similar reductions in mean fibril height (59 nm) and width (55 nm) suggest isometric shrinkage of the fiber cross section during contraction. SFs feature a complex top surface possibly because of associated regulatory proteins or organization of the fibers into structurally distinct functional sarcomeric units. However, despite the significant reduction in fibril thickness during contraction, local roughness (arithmetic mean roughness, *Ra*) and global roughness (waviness *Wq*) values along longitudinal cross sections before and after contraction were similar (*Ra* 19.2 nm and 22.04 nm, *Wq* 512 nm and 472nm, respectively), indicating preservation of these structural variations despite the overall drop in fibril height after ATP exposure (Fig. 4G).

**Figure 4.**
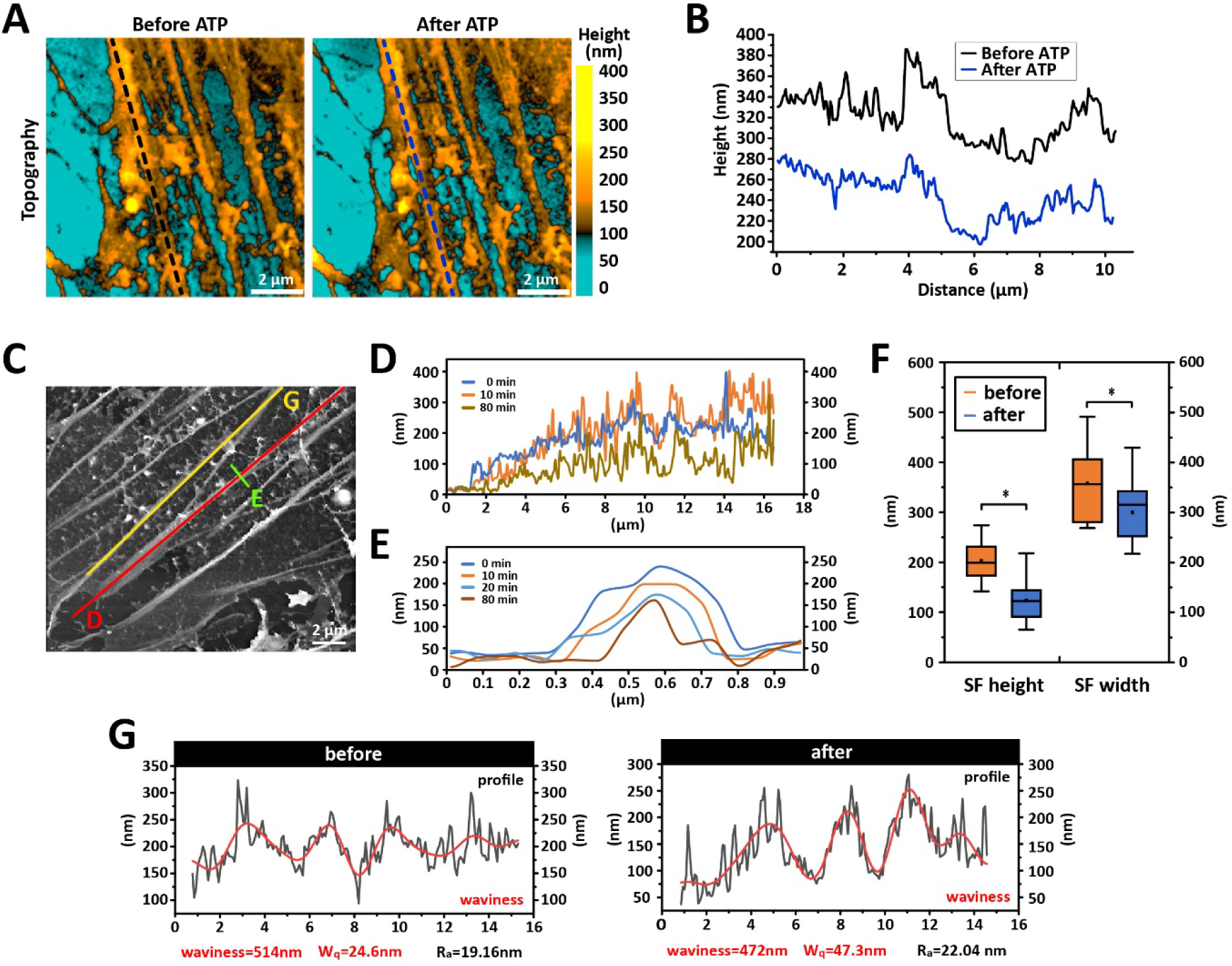
**(A)** SICM topographic image from a timelapse series depicting ATP-induced contracting SFs in a de-roofed cell sample. **(B)** Height profiles generated along the dashes lines in **(A)** before ATP (black) and after (blue) ATP stimulated SF contraction. **(C)** Representative SICM topography of exposed SFs in a de-roofed cell. Colored lines correspond to height cross-sections displayed in **(D)**, **(E)**, and **(G)**. **(D)** Longitudinal SF height profiles generated along the red line indicated in **(C)** at 0, 10, and 80 min timepoints after first ATP addition. **(E)** Orthogonal SF height profiles generated along the green line indicated in **(C)** at different time points after ATP addition. **(F)** Box plots (line: median, dot: mean and box: 25/75% range, whiskers: min/max range) depicting the reduction in SF height and width after ATP-induced contraction. The * symbol indicates statistical significance (p-values <0.05) according to a paired t-test, N=15. **(G)** Long-range (waviness, red line) and local topography (roughness, black line) changes determined along the yellow profile line shown in **(C)**.

The height reduction in some fibrils was accompanied by the formation of longitudinal grooves extending several hundred nm in length and reaching a depth of ∼100 nm (Fig. 5A and 5B). The apparent splitting of the SF into two separate cable-like structures at the fibril edge suggests the presence of distinct functional units with higher resistance to ATP-induced morphological changes compared to the central region of the fibril, as has been previously proposed^26^.

**Figure 5.**
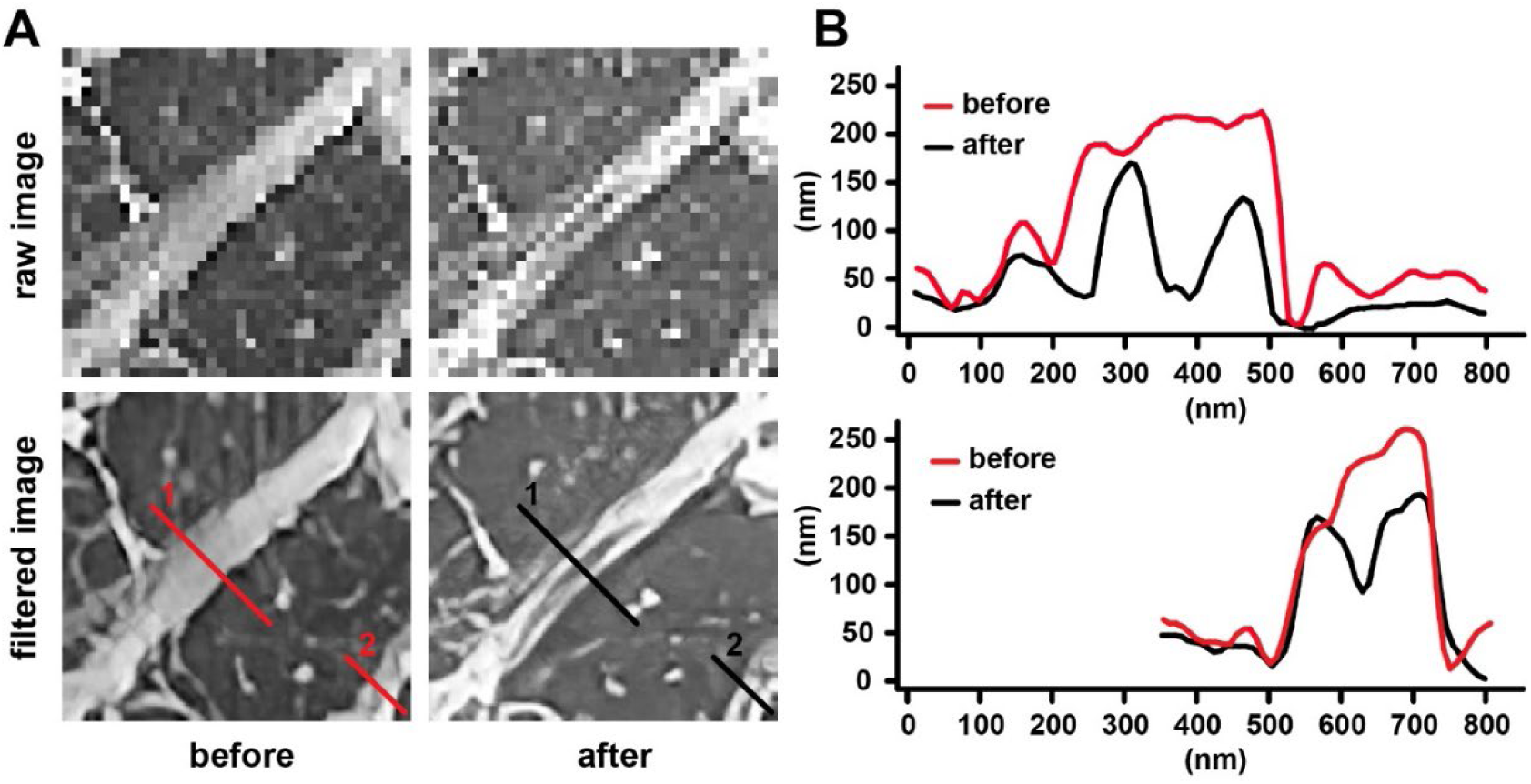
**(A)** Raw (upper panels) and filtered (lower panels) SICM topography images before and after ATP stimulated SF contraction. **(B)** Height profiles extracted at the positions indicated in **(A)**.

### Characterization of mechanical changes during SF contraction

In addition to its imaging capability, SICM is a valuable tool for characterizing mechanical properties of biological samples, including living cells^17,39,40^, with submicrometer lateral resolution^41^. During SICM nanomechanics measurements, local sample stiffness is assessed by the ability to resist deformation forces exerted by the SICM tip at elevated ion current setpoints. Compared to AFM, SICM can detect significantly smaller sample deformations and requires lower indentation forces^40,41^, making it especially suitable for the characterization of soft, and/or thin samples that cannot be indented deeply. An additional advantage of SICM is the ability to record topographic images and sample deformation maps simultaneously^40^. Here we used SICM deformation mapping to characterize mechanical changes in the de-roofed cell sample after ATP-induced SF contraction. For this, we first collected topographic images of an exposed SF array before ATP addition at ion-current reduction setpoints of 0.5%, 1%, and 2% (Fig. 6A). Sample deformation was negligible on the cell-free glass substrate (0 ± 4 nm), intermediate on SF-free membrane areas (29 ± 26 nm), and highest on SF-containing regions (74 ± 42 nm). Collecting a second set of topographic maps on the same sample area after ATP stimulation showed decreased deformation in SF-containing regions (61 ± 48 nm), accompanied by a shift of the deformation distribution toward lower values (Fig. 6A,B). In contrast, deformation remained unchanged in SF-free membrane regions (30 ± 31 nm) and on the bare glass substrate (1 ± 1 nm). Reduced sample deformation exclusively in SF-containing regions indicated selective SF stiffening after ATP stimulation. Plotting a differential deformation map (subtracting deformation maps collected before and after ATP addition) of a representative SF furthermore showed that the scale of ATP-induced stiffening varied along the SF, consistent with the presence of SF segments with different mechanical properties after ATP-induced stiffening (Fig. 6C).

**Figure 6.**
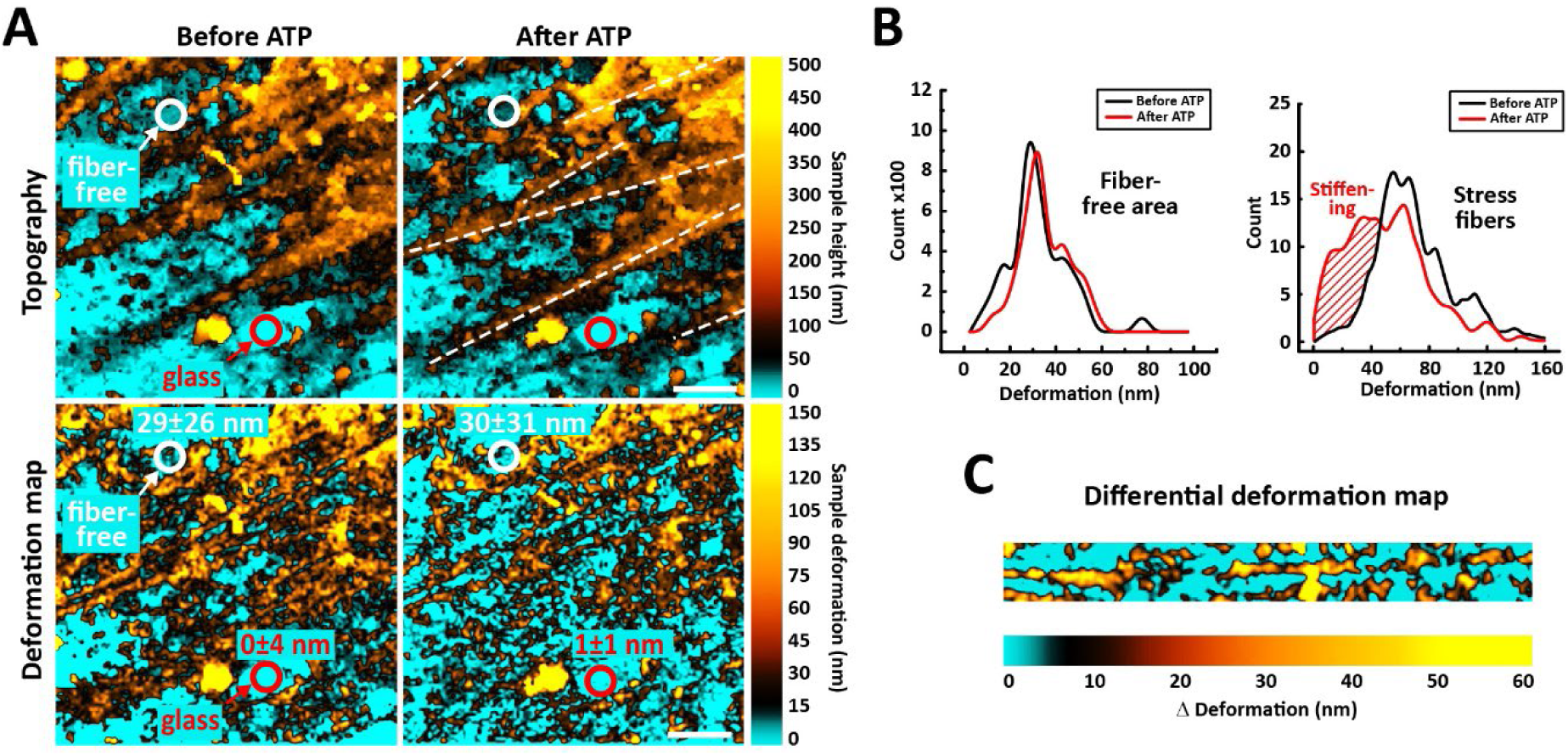
Nanomechanical characterization of contracting SFs. **(A)** SICM topographies (upper panels) and deformation maps (lower panels) of SFs in a deroofed cell sample before and after ATP addition. White circles denote a representative fiber-free cell membrane area, red circles denote an area of the glass substrate, and corresponding deformation values (mean ± SD). Scale bar 1.5 µm. **(B)** Deformation distributions before and after ATP stimulation of SFs (right) and fiber-free areas (left). SF deformation values were collected along the dashed lines shown in **(A)**. **(C)** SF differential deformation map of an individual SF undergoing inhomogeneous fiber stiffening after ATP exposure.

## Discussion

In this study we combined cell de-roofing and SICM imaging to investigate dynamic structural and mechanical changes during actin SF contraction. The principal advantage of this approach is that de-roofing provides direct nanoprobe access to SFs while preserving their native contractile state, allowing for high-resolution imaging and nanomechanical characterization (Figure 7). This approach differs from previous SPM studies, which investigated cytoskeletal structures indirectly across the overlying plasma membrane. Indeed, AFM images of living adherent cells often display prominent SFs and other cortical actin structures because the compliant plasma membrane is displaced against these stiffer cytoskeletal elements by the scanning AFM probe. This phenomenon has been exploited to visualize dynamic cortical actin rearrangements using both conventional^42^ and high-speed AFM^43,44^, while mechanical properties of actin SFs in living cells have been deduced using conventional^45^ or torsional mode AFM^46^.

**Figure 7.**
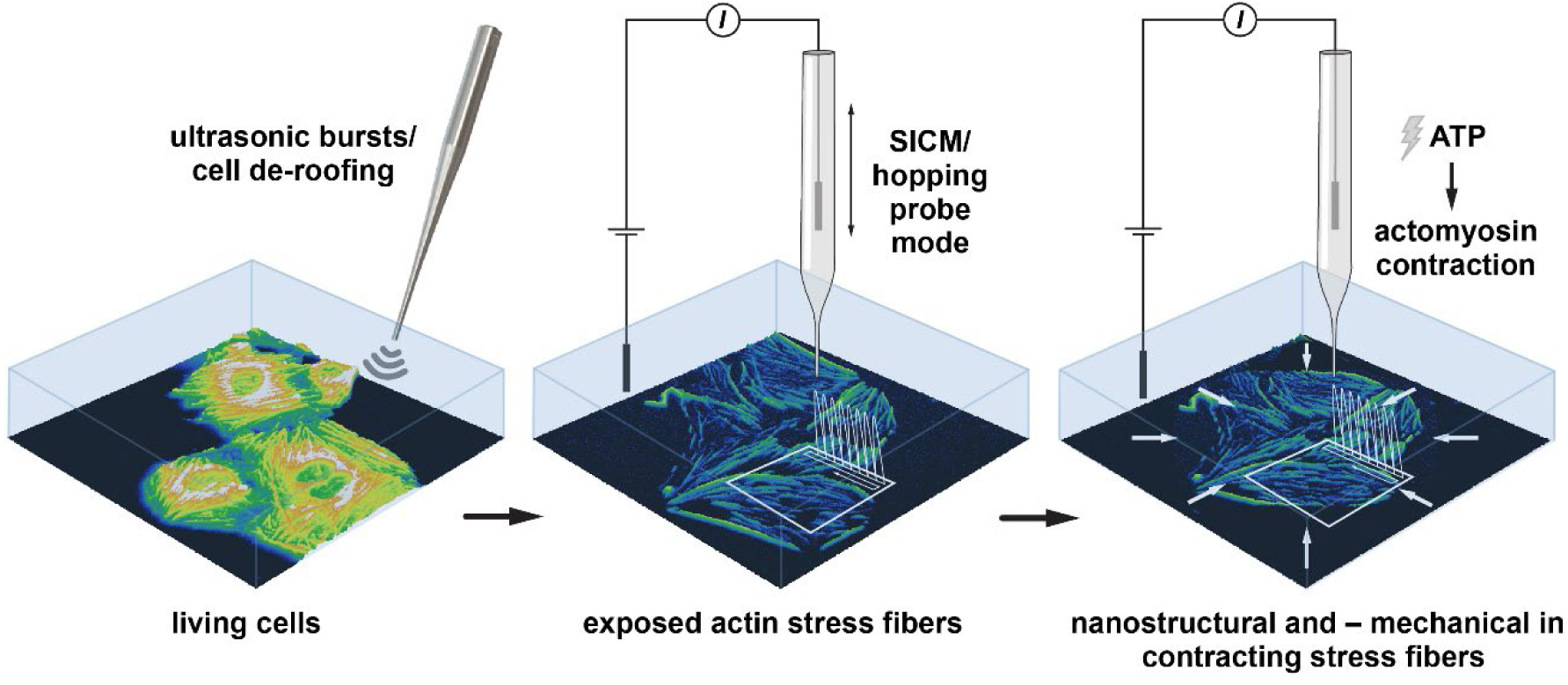
Integrating ultrasonication-based cell de-roofing and SICM for the structural and mechanical characterization of contracting actin SFs.

In contrast to cell de-roofing experiments, live-cell investigations permit the real-time visualization of physiological actomyosin-driven SF remodeling. However, the overlying plasma membrane and the associated apical actin cortex may obscure nanoscale features on the surface of ventral SFs and complicate the interpretation of nanomechanical measurements. While exposed SFs can also be investigated by AFM, hopping mode SICM offers distinct advantages owing to its non-contact imaging modality, which minimizes lateral probe forces and reduces sample distortion. In addition, the high-aspect-ratio of the nanopipette apex provides reliable imaging of steep sample edges which is often challenging for AFM because of limitations in scanner z-range and tip geometry. SICM is therefore particularly well suited for imaging exposed SFs and other intracellular structures with abrupt, steep changes in height, and may cause less damage to complex samples than an AFM tip.

The SICM images of resting and contracting SFs not only corroborated previous observations of SF structure and mechanics but also revealed novel nanoscale features that could not be observed by other imaging methods. In particular, the excellent resolution z-direction (∼10 nm) of SICM enabled us to measure the height of native resting and contracting SFs with high accuracy, while its lateral resolution (∼50 nm) exceeded that of conventional and some super-resolution optical microscopy variants. The excellent resolution in Z-direction of SICM revealed that SFs feature a complex nano-topography characterized by long-range height modulations (period ∼500 nm, height ∼25 nm) overlayed with irregular corrugations (R_a_ ∼19.2 nm). EM images of extracted SFs suggest a helical arrangement of embedded microfilament subbundles^47^, and the observed long-range waviness may reflect the helix pitch of such structures. Likewise, the observed contraction-induced SF splitting corresponds to the separation of individual fiber sub-bundles, as previously proposed based on AFM analysis^26^.

Compared to nanometric AFM tips, SICM nanopipettes generally provide lower lateral resolution because both pipette aperture and side wall geometry contribute to sample convolution^48^. Nevertheless, recent advances in fabricating nanopipettes with small pore diameters of <10 nm^49,50^ and the implementation of advanced image deconvolution methods have contributed to enhancing the lateral resolution of SICM^51^. To ensure accurate sample contouring, we used comparatively low frame rates (2 to 5 min per frame). However, remaining scan speed disadvantages of SICM are being progressively alleviated through the implementation of dedicated high-speed SICM scanners^52–54^ and computationally-driven optimization of scan line data acquisition^55^. These advances should further expand the utility of SICM for high-resolution, real-time imaging of intracellular structures and dynamics.

Microsonication-based cell de-roofing is particularly effective for exposing intracellular structures tightly associated with the basal cell membrane, including the actin cortex^23^ and clathrin-coated pits^25^, whereas softer or larger intracellular structures and organelles are typically removed during the procedure. The continuing development of gentler cell de-roofing methods that selectively remove the plasma membrane while preserving fragile intracellular structures could substantially broaden the scope of SICM-based intracellular investigation. Although recent advances in AFM tip design have produced long and ultrathin tips capable of transient membrane penetration for intracellular exploration without compromising cell viability^56 57^, long and ultrathin tips have become available that permit transient membrane penetration for intracellular exploration without compromising cell viability^56^. However, force curve-based intracellular exploration by SPM remains limited in spatial and temporal resolution. Consequently, high-resolution imaging of dynamic intracellular currently still relies on prior cell membrane removal, highlighting the continued importance of de-roofing approaches for intracellular nanoscale imaging.

The SICM topographs confirmed the characteristic splayed arrangement of individual actin filaments at FAs, in agreement with previous EM^6,58^ and AFM studies of fixed de-roofed cells^22^. While actin filaments are bundled into approximately cylindrical SFs, the flattening and spreading of the filament array at SF termini may maximize their engagement with membrane-proximal integrin-associated linker complexes. Notably, timelapse SICM imaging revealed that this frayed, flattened FA architecture is largely retained during SF retraction and FA sliding, suggesting that efficient force transmission and integrin-mediated anchorage are preserved throughout contraction-induced SF shortening.

Timelapse SICM imaging furthermore revealed an intricate network of crosslinks between neighboring SFs that persisted throughout ATP-induced contraction, demonstrating coordinated mechanical coupling between adjacent fibers and their integration into a broader cytoskeletal network. Through associated actomyosin filaments, SFs connect to the cortical actin mesh, forming a continuous, contractile network^59–61^. Moreover, SFs closely associate with vimentin intermediate filament networks^62^, and vimentin integration regulates SF contraction^63^. In addition, spectrin networks stabilize SFs and anchor them to the plasma membrane^64^. As a label-free technique, SICM can visualize these intertwined contractile networks simultaneously under native conditions. However, SICM topographs permit no direct molecular identification, but specific fluorescent labeling and complementary optical superresolution microscopy could help to identify individual cytoskeletal components in the high-resolution SICM images.

Previous experiments using extracted native, free-floating SFs showed strong contraction of up to 23% in response to ATP^47^. In contrast, in our assay SFs remained attached to the substrate and contracted considerably less. Although contraction affected primarily the distal SF ends, the SF diameter nevertheless decreased significantly even in central SF regions that displayed no translocation, probably because of tighter packing of actin filaments^47^. SF also stiffened after ATP stimulation, indicated by increasing deformation resistance to the hydrodynamic pressure applied by the nanopipette at elevated approach setpoints. When used in combination with suitable calibration substrates with well-known elastic properties (for instance decane droplets), SICM can quantitatively analyze nanomechanical properties of living cells^40^. However, these experiments typically employ nanopipettes with larger tip diameters (∼100 nm), whereas we used smaller nanopipettes (∼50 nm diameter) so both high-resolution imaging and subsequent nanomechanical characterization of individual SFs could be performed with a single nanopipette. Quantitative contact models that describe the dependence of sample stress values on nanopipette tip diameters <80 nm have not been established, and we therefore limited our analysis to a qualitative assessment of SF stiffening based on changes in sample deformation. Nevertheless, our approach revealed substantial spatial variation in stiffening along individual SFs during contraction. Comparable local differences in contraction-induced SF stiffening could be related to the sarcomeric organization of non-muscle myosin II inside SFs and have been reported in living cells using AFM nanoindentation measurements^65^, supporting the notion that the de-roofing procedure preserves the native mechanical behavior of SFs and demonstrating that SICM can effectively resolve such heterogeneity.

In conclusion, we demonstrate that SICM combined with microsonication-based cell de-roofing enables direct, high-resolution structural and mechanical imaging of intracellular actin architectures under near-physiological conditions. This approach revealed nanoscale features of SF organization, contraction-associated remodeling, and spatially heterogeneous mechanical responses that are difficult to access with existing imaging techniques. Beyond investigating actin SFs, the integration of SICM with controlled cell membrane removal, specific molecular labeling, and correlative imaging approaches offers a versatile framework for investigating the structure–mechanics relationships of diverse intracellular assemblies. Improvements in nanopipette fabrication, scanning speed, and quantitative nanomechanics modelling continue and will further expand the spatial and temporal resolution of intracellular SICM imaging, positioning SICM as an important tool for studying intracellular organization and dynamics at nanometer resolution.

## Methods

### Cell culture, cell de-roofing, and holographic tomography imaging

U2OS cells stably expressing ß-actin-tagGFP (U2OS-actin-GFP, CellTrend GmbH) were maintained in DMEM 10% FCS and 100U penicillin/100mg*ml^−1^. For de-roofing, cells were seeded onto 35mm glass bottom (FluoroDish, WPI) or plastic cell culture dishes and grown for 48 h to about 75% confluency. After two rinses with phosphate-buffered saline (PBS), cells were incubated for 60 sec in 1 mM PLL/PBS (MW 15-30kD, Sigma Aldrich), rinsed two times in ice-cold cell de-roofing buffer (30 mM KCl, 5 mM MgCl2, 10 µM CaCl2, 10 mM Tris pH 7.0, 4% PEG) containing protease inhibitors (cOmplete EDTA-free Mini Protease Inhibitor Cocktail, Roche) and de-roofed using several pulses from a handheld microsonicator (UP50H, Hielscher Ultrasound Technology) on its lowest power setting. After sonication cells were washed 5 times in de-roofing buffer without PEG and protease inhibitors and directly imaged by SICM or fluorescence microscopy (U2OS-actin-GFP). Living cells were transferred into CO_2_-independent cell culture medium (ThermoFisher) for SICM imaging. For holographic tomography imaging, de-roofed cells seeded in 35mm glass bottom dishes were maintained in 1 ml of de-roofing buffer (without inhibitors and PEG) and imaged at RT before and after addition of 1 mM ATP/Mg^2+^ using a 3D Explorer (NanoLive) with fluorescence imaging capability.

### Fabrication of SICM glass nanopipettes

SICM glass nanopipettes for high-resolution scanning and mechanical measurements were fabricated by pulling borosilicate glass capillaries (inner diameter = 0.58 mm, outer diameter = 1.00 mm) into nanopipettes with a tip diameter of ∼50 nm using a CO_2_ laser puller (Model P-2000, Sutter Instruments Co., USA) using a preheated two-step protocol as described previously^31^. Briefly, nanopipettes were fabricated from borosilicate glass capillaries using a two-step pulling protocol. Prior to pulling, capillaries were preheated for 5 s (heat 370, filament 3, velocity 25, delay 150, pull 0) to optimize the inner and outer tip geometry. The first pulling step was performed using heat 310, filament 3, velocity 25, and delay 150, followed by a second step using heat 280, filament 2, velocity 23, delay 150, and pull 250. Before each SICM experiment, the nanopipettes were routinely checked by measuring their electrical resistance, which was ∼300 MΩ when filled with PBS.

### Combined SICM and fluorescence microscopy setup

For simultaneously obtaining fluorescence and topographic images of exposed SFs, a previously described SICM scanner system^52,66^ was mounted on an optical microscope ECLIPSE TiU (Nikon, Japan), in turn resting on an antivibration Herz TS-150 table (Herzan, USA). Briefly, a glass-bottom dish with the de-roofed cell sample (in cell de-roofing buffer) was placed onto the optical microscope and the nanopipette was backfilled with imaging solution (de-roofing buffer). An Ag/AgCl electrode was then inserted into the nanopipette and another Ag/AgCl reference electrode placed into the cell culture dish for measuring the ion current flow through the nanopipette using a homemade 1 GΩ feedback resistance current amplifier at an applied bias potential of 200 mV. The nanopipette probe was initially approached onto the sample using a stepping motor (KXC06020-GC, SURUGA SEIKI, Japan), and a homemade *Z*-piezo stage for feedback-controlled final sample approach and scanning. The precise nanoprobe position was controlled with a 25 x 25 µm^2^ *XY*-piezo stage, while a manually operated manipulator (BSS76−60C, SURUGA SEIKI, Japan) with a travel range of ± 6.5 mm allowed for additional scan frame *XY*-positioning. Signal collection and scan control were performed using custom-developed software in LabVIEW2014 (National Instruments, USA).

### Image processing

Fluorescence microscopy images were processed in ImageJ and Adobe Photoshop. SF translocation was visualized using the iterative particle velocimetry (PIV) module in ImageJ. When indicated, SICM raw images were filtered using Topaz Sharpen AI (Topaz Labs).

### Measuring mechanical properties of SFs by SICM

Combined SICM scanning and mechanical mapping was performed using a customized SICM setup operated in adaptive resolution hopping probe HPICM mode^16^ using an ICAPPIC Controller (IC-UN-001, ICAPPIC Ltd, UK) as previously described^40,67^. Briefly, the SICM scan head consisted of a PIHera P-621.2 XY Nanopositioning Stage (Physik Instrumente, Germany) with a 50 × 50 µm travel range for moving the sample and a LISA piezo actuator P-753.21 C (Physik Instrumente, Germany) with a travel range of 25 µm for pipette positioning along the Z-axis. The SICM scan head was mounted on an inverted optical Eclipse Ti-2 microscope (Nikon, Japan) and covered by a Faraday cage for electrical noise shielding. The SICM control, data acquisition, and analysis software were written and provided by Dr Pavel Novak, ICAPPIC Ltd. Ion currents were monitored using a MultiClamp 700B amplifier (Molecular Devices, USA) and a 2 kHz low-pass filter. A typical external holding voltage of 200 mV was supplied to the scanning nanopipette. For SF deformation mapping during SICM scanning, the nanopipette approach rate was set to 200 μm/s, and images were acquired using a three-setpoint mode, as previously described by Kolmogorov et al.^40^: First, a non-contact topographic image (256 x 256 pixel) was recorded at an ion current decrease of 0.5%. Subsequently, two further topographic images were collected at ion current decreases of 1% and 2%, inducing increasing sample deformation due to hydrodynamic pressure applied by the nanopipette at these elevated setpoints. Setpoint-dependent sample deformations maps were determined from the apparent height difference between the 1% and 2% setpoint images using SICM ImageViewer (ICAPPIC Ltd., London, UK).

## Acknowledgements

This work was supported the World Premier International Research Center Initiative (WPI), MEXT, Japan. Y.Z and Y.K. were supported by Japan Society for the Promotion of Science KAKENHI 21H01770, 22K04890, and 23K21070. C.M.F. received support from Japan Society for the Promotion of Science KAKENHI 20H03218. Y.-R.L. received support from the Japan-Taiwan Exchange Association. This research was also supported by AMED under Grant Number JP21wm0525015, the JST FOREST Program (Grant Number JPMJFR203K), the Hibi Science Foundation, a Grant-in-Aid for Scientific Research (S) (23H05480), AMED-CREST (JP23gm1410012), CREST (JPMJCR24T6) from the Japan Science and Technology Agency (JST), and the Human Frontier Science Program (HFSP) (no. RGP0028/2022).

## Author contributions

Conceptualization: Y.Z., C.M.F.; Methodology: Y.Z., Y.T., Y.-R.L., A.S., Y.K., C.M.F.; Validation: Y.Z., Y.T., Y.K., C.M.F; Formal analysis: Y.Z., Y.T., Y.K., C.M.F; Investigation: Y.Z., C.M.F.; Resources: Y.Z., C.M.F.; Writing - original draft: Y.Z, C.M.F.; Writing - review & editing: Y.Z., Y.T., Y.K., Y.-R.L., C.M.F.; Visualization: Y.Z., Y.T., Y.K., C.M.F; Project administration: Y.Z., C.M.F.; Funding acquisition: Y.Z, Y.K., Y.-R.L., Y.T., C.M.F.

## Supplementary Files

**Supplementary Figure S1.** SICM image of a living (A) and an unfixed, partially de-roofed U2OS cells (B).

**Supplementary Figure S2.** SICM topographical images of stress fibers (SFs) acquired immediately after de-roofing (A) and of the same SFs after 2 h of continuous scanning at room temperature in fully de-roofed U2OS cells (B).

**Supplementary Movie 1. Stress fibers remain contractile after cell de-roofing.** De-roofed U2OS cells expressing tagGFP-actin were simultaneously imaged by fluorescence (left) and 3D holotomography (middle) before (top) and after stimulation with 1 mM ATP/Mg2+. The right image represents a twofold magnified view of a subregion of the 3D holotomography movie. The uncropped image size corresponds to 90 x 90 µm2.

**Supplementary Movie 2. Timelapse SICM image series of SFs contracting in response to ATP stimulation.** ATP/Mg^2+^ complex was added in individual doses of 1 mM up to a total final concentration of 4 mM. The full range of the height scale corresponds to 2 μm.

**Supplementary Movie 3. SFs contract as mechanically-coupled networks.** The frames of the SICM timelapse series shown in Movie 2 (frame acquisition time 10 min) were corrected for *x,y*-drift, and SFs and associated interconnections were traced in red using the ridge detection function in ImageJ.

