## Supplemental Information for "Investigating nanostructure and -mechanics of contracting actin stress fibers by scanning ion conductance microscopy"

**Supplementary Figure S1**

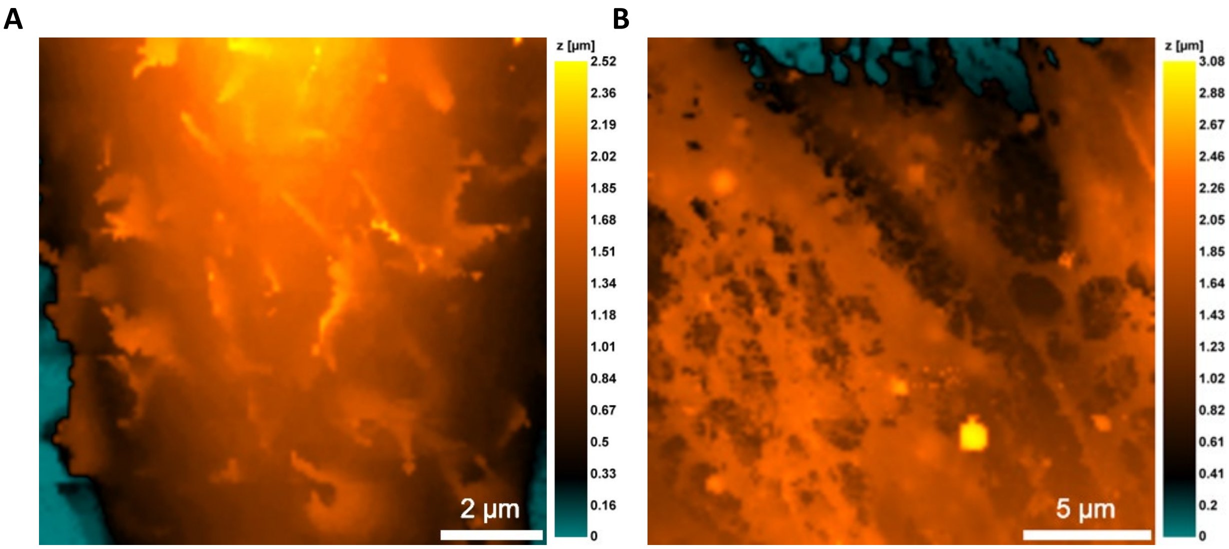


**Supplementary Figure S1**. SICM topographical images of a living (A) and an unfixed, partially de-roofed U2OS cell (B).

**Supplementary Figure S2**

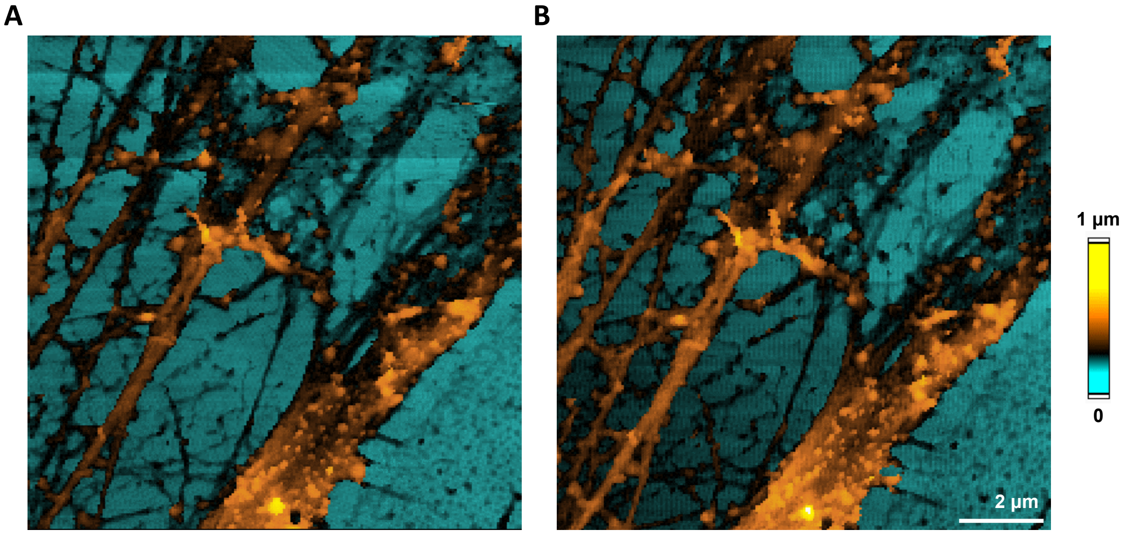


**Supplementary Figure S2.** SICM topographical images of stress fibers (SFs) acquired immediately after de-roofing (A) and of the same SFs after 2 h of continuous scanning at room temperature in fully de-roofed U2OS cells (B).

**Supplementary Movie 1. Stress fibers remain contractile after cell de-roofing.** De-roofed U2OS cells expressing tagGFP-actin were simultaneously imaged by fluorescence (left) and 3D holotomography (middle) before (top) and after stimulation with 1 mM ATP/$Mg^{2+}$. The right image represents a twofold magnified view of a subregion of the 3D holotomography movie. The uncropped image size corresponds to 90 x 90 $\mu m^2$.

**Supplementary Movie 2. Timelapse SICM image series of SFs contracting in response to ATP stimulation.** ATP/$Mg^{2+}$ complex was added in individual doses of 1 mM up to a total final concentration of 4 mM. The full range of the height scale corresponds to 2 μm.

**Supplementary Movie 3. SFs contract as mechanically-coupled networks.** The frames of the SICM timelapse series shown in Movie 2 (frame acquisition time 10 min) were corrected for *x,y*-drift, and SFs and associated interconnections were traced in red using the ridge detection function in ImageJ.
